# Nirmatrelvir, Molnupiravir, and Remdesivir maintain potent *in vitro* activity against the SARS-CoV-2 Omicron variant

**DOI:** 10.1101/2022.01.17.476685

**Authors:** Romel Rosales, Briana L. McGovern, M. Luis Rodriguez, Devendra K. Rai, Rhonda D. Cardin, Annaliesa S. Anderson, PSP study group, Emilia Mia Sordillo, Harm van Bakel, Viviana Simon, Adolfo García-Sastre, Kris M. White

**Author notes:** PSP (Pathogen Surveillance Program) study group: Hala Alshammary, R. Banu, K. Farrugia, Ana Silvia Gonzalez-Reiche, A. Paniz-Mondolfi, J. Polanco.

## Abstract

Variants of SARS-CoV-2 have become a major public health concern due to increased transmissibility, and escape from natural immunity, vaccine protection, and monoclonal antibody therapeutics. The highly transmissible Omicron variant has up to 32 mutations within the spike protein, many more than previous variants, heightening these concerns of immune escape. There are now multiple antiviral therapeutics that have received approval for emergency use by the FDA and target both the SARS-CoV-2 RNA-dependent RNA polymerase (RdRp) and the main protease (Mpro), which have accumulated fewer mutations in known SARS-CoV-2 variants. Here we test nirmatrelvir (PF-07321332), and other clinically relevant SARS-CoV-2 antivirals, against a panel of SARS-CoV-2 variants, including the novel Omicron variant, in live-virus antiviral assays. We confirm that nirmatrelvir and other clinically relevant antivirals all maintain activity against all variants tested, including Omicron.

## Introduction

Coronavirus disease 2019 (COVID-19) has caused a global pandemic and millions of deaths due to the viral agent, SARS-CoV-2. The catastrophic spread of COVID-19 has severely impacted lives, which continues with the introduction of new variants, across the globe and has been declared a public health emergency of international concern^1^.

Coronaviruses such as SARS-CoV-2 undergo genetic drift during replication in host organisms through mutations generated by an error-prone RNA-dependent RNA polymerase (RdRp).

Populations of SARS-CoV-2 that inherit the same distinctive mutations are termed variants. Over the past year, the emergence of new variants has become a major factor in the ongoing COVID-19 pandemic. Many variants of SARS-CoV-2 have been discovered; however, only some such as the Delta and Omicron variants have recently been identified and labeled as variants of concern (VOC) because of increased transmissibility and evasion of natural and vaccine-induced immune responses^2,3^.

The majority of mutations observed in the novel Omicron variant occur within the spike protein, likely driven by the dominant nature of spike protein epitopes in the immune response to SARS-CoV-2 infection and vaccination^4^. The Omicron variant is characterized by a large number of mutations, with 26 to 32 changes in the spike protein^5^. While this is of major concern to the current SARS-CoV-2 vaccines and monoclonal antibody therapeutics^6–8^, that exclusively target the spike protein, the more recently authorized antiviral therapeutics for the treatment of COVID-19 target other viral proteins which have accumulated fewer mutations throughout the emergence of SARS-CoV-2 variants. Nirmatrelvir (the SARS-CoV-2 antiviral component of Paxlovid) targets the main protease (Mpro or 3CL protease) of SARS-CoV-2 and has demonstrated nearly a 90% reduction in hospitalization and death in clinical trials^9,10^. Molnupiravir and remdesivir target the viral RdRp and have shown mixed efficacy in the clinic^11,12^. It is crucial to evaluate the antiviral efficacy of these clinically effective therapeutic antivirals that will play a critical role in the public health response to current and future SARS-CoV-2 variants. Here we assess the susceptibility of the novel Omicron variant of SARS-CoV-2 to the important clinical antiviral nirmatrelvir, compared to other clinically relevant antiviral therapeutics.

## Materials and Methods

Two thousand Vero-TMPRSS2 or HeLa-ACE2 cells (BPS Bioscience) were seeded into 96-well plates in DMEM (10% FBS) and incubated for 24 hours at 37°C, 5% CO_2_. Two hours before infection, the medium was replaced with 100 μL of DMEM (2% FBS) containing the compound of interest at concentrations 50% greater than those indicated, including a DMSO control. Plates were then transferred into the BSL3 facility and 100 PFU (MOI = 0.025) was added in 50 μL of DMEM (2% FBS), bringing the final compound concentration to those indicated. Plates were then incubated for 48 hours at 37°C. After infection, supernatants were removed and cells were fixed with 4% formaldehyde for 24 hours prior to being removed from the BSL3 facility. The cells were then immunostained for the viral N protein (an inhouse mAb 1C7, provided by Dr. Thomas Moran) with a DAPI counterstain. Infected cells (488 nm) and total cells (DAPI) were quantified using the Celigo (Nexcelcom) imaging cytometer. Infectivity was measured by the accumulation of viral N protein (fluorescence accumulation). Percent infection was quantified as ((Infected cells/Total cells) - Background) *100 and the DMSO control was then set to 100% infection for analysis. Data was fit using nonlinear regression and IC50s for each experiment were determined using GraphPad Prism version 8.0.0 (San Diego, CA). Cytotoxicity was also performed using the MTT assay (Roche), according to the manufacturer’s instructions. Cytotoxicity was performed in uninfected cells with same compound dilutions and concurrent with viral replication assay. All assays were performed in biologically independent triplicates. Vero-TMPRSS2 experiments were performed in the presence of the Pgp-inhibitor CP-100356 (2 μM) in order to limit compound efflux.

Nasopharyngeal swab specimens were collected as part of the routine SARS-CoV-2 surveillance conducted by the Mount Sinai Pathogen Surveillance program (IRB approved, HS#13-00981). Specimens were selected for viral culture on Vero-TMPRSS2 (BPS Bioscience) cells based on the complete viral genome sequence information^13^. The SARS-CoV-2 virus USA-WA1/2020 was obtained from BEI resources (NR-52281) and used as wild-type reference. Viruses were grown in Vero-TMPRSS2 cells (BPS Bioscience) for 4–6 d; the supernatant was clarified by centrifugation at 4,000*g* for 5 min and aliquots were frozen at −80°C for long term use. Expanded viral stocks were sequence-verified to be the identified SARS-CoV-2 variant and titered on Vero-TMPRSS2 cells before use in antiviral assays.

## Results and Discussion

We have established live-virus immunofluorescence-based antiviral assays with many relevant variants of SARS-CoV-2, including the novel Omicron variant, in both Vero-TMPRSS2 and HeLa-ACE2 cells. Six-point dose response curves were generated for nirmatrelvir antiviral activity against the SARS-CoV-2/WA1 strain, a previously described mouse adapted SARS-CoV-2 strain (MA-SARS-CoV-2)^14^, and representative Alpha, Beta, Delta, and Omicron variants. Variants were collected from deidentified nasopharyngeal swab specimens as part of the routine SARS-CoV-2 surveillance conducted by the Mount Sinai Pathogen Surveillance program. Cytotoxicity assays were run concurrently with the antiviral assays in uninfected cells using matched conditions.

We found that the potency of nirmatrelvir was similar across all variants tested (Fig. 1+2 and Table 1), including Omicron, as compared to the SARS-CoV-2/WA1 strain in both Vero-TMPRSS2 and HeLa-ACE2 cells. EIDD-1931 (the active metabolite of molnupiravir) and remdesivir also maintained activity across variants. Vero-TMPRSS2 experiments were performed in the presence of the Pgp-inhibitor CP-100356 (2 μM) in order to limit compound efflux (Table 2). These results are congruent with the relative conservation observed in the SARS-CoV-2 Mpro and RdRp, as compared to the spike protein, across all variants.

**Figure 1.**
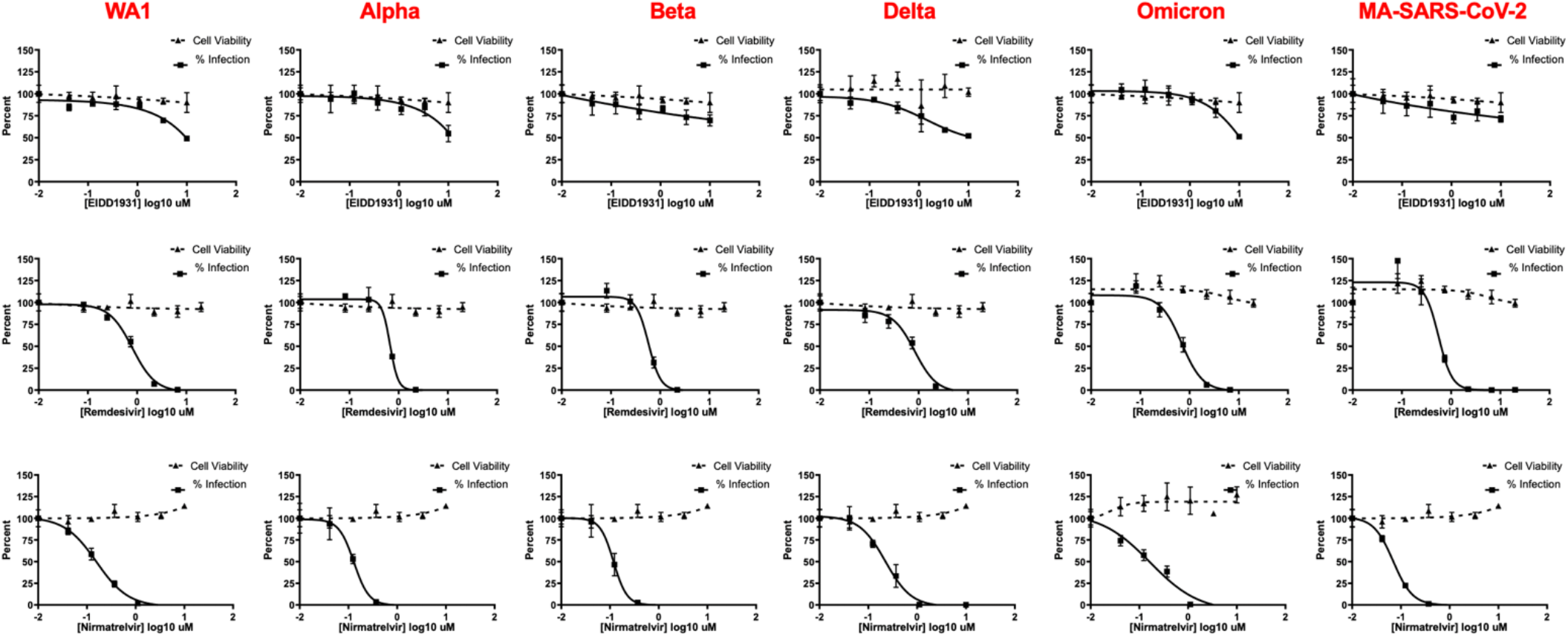
IF-based live-virus antiviral (solid lines) and MTT-based cytotoxicity (dashed lines) dose response curves for nirmatrelvir, remdesivir, and EIDD-1931 (molnupiravir) against a panel of SARS-CoV-2 variants in HeLa-ACE2 cells. The IC_50_ and CC_50_ of each compound against each variant is indicated in Figure 2 and Table 1. No loss of activity was observed for tested inhibitors against the Omicron variant. Data are means ± SD of a representative experiment performed in biological triplicate.

**Figure 2.**
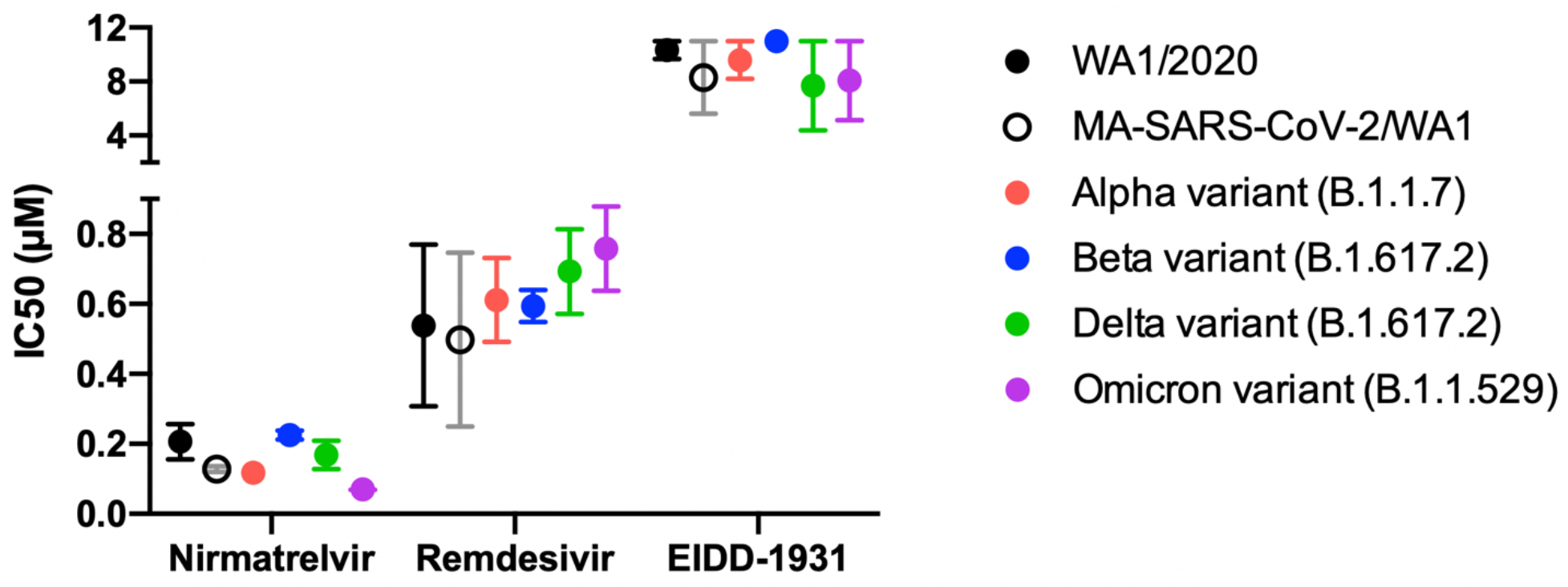
Antiviral IC_50_ calculated from 6-point dose-response curves in IF-based live-virus antiviral assays for nirmatrelvir, remdesivir, and EIDD-1931 (molnupiravir) against a panel of SARS-CoV-2 variants in HeLa-ACE2 cells. The IC_50_ was calculated and graphed using GraphPad Prism version 8.0.0. No loss of activity was observed for tested inhibitors against the Omicron variant. Data are means ± SD of two independent experiments performed in biological triplicate.

**Table 1.** Nirmatrelvir IC_50_, IC_90_, and CC_50_ against clinically relevant SARS-CoV-2 variants in HeLa-ACE2 cells (n=6).

|  | Nirmatrelvir |  |  | Remdesivir |  |  | EIDD-1931 |  |  |
| --- | --- | --- | --- | --- | --- | --- | --- | --- | --- |
| Variant | IC<sub>50</sub><br>(μM) | IC<sub>90</sub><br>(μM) | CC<sub>50</sub><br>(μM) | IC<sub>50</sub><br>(μM) | IC<sub>90</sub><br>(μM) | CC<sub>50</sub><br>(μM) | IC<sub>50</sub><br>(μM) | IC<sub>90</sub><br>(μM) | CC<sub>50</sub><br>(μM) |
| SARS-CoV-2 USA-WA1/2020 | 0.21 | 0.72 | >10 | 0.54 | 2.34 | >20 | 10.33 | 26.61 | >10 |
| (mouse-adapted) MA-SARS-CoV-2/WA1 | 0.13 | 0.27 | >10 | 0.50 | 1.20 | >20 | 8.30 | >10 | >10 |
| SARS-CoV-2 Alpha variant (B.1.1.7) | 0.12 | 0.24 | >10 | 0.61 | 1.02 | >20 | 9.6 | >10 | >10 |
| SARS-CoV-2 Beta variant (B.1.617.2) | 0.23 | 0.78 | >10 | 0.60 | 1.09 | >20 | >10 | >10 | >10 |
| SARS-CoV-2 Delta variant (B.1.617.2) | 0.17 | 1.17 | >10 | 0.69 | 2.16 | >20 | 12.79 | 7.7 | >10 |
| SARS-CoV-2 Omicron variant (B.1.1.529) | 0.07 | 0.19 | >10 | 0.76 | 2.01 | >20 | 10.44 | 8.1 | >10 |

**Table 2.**
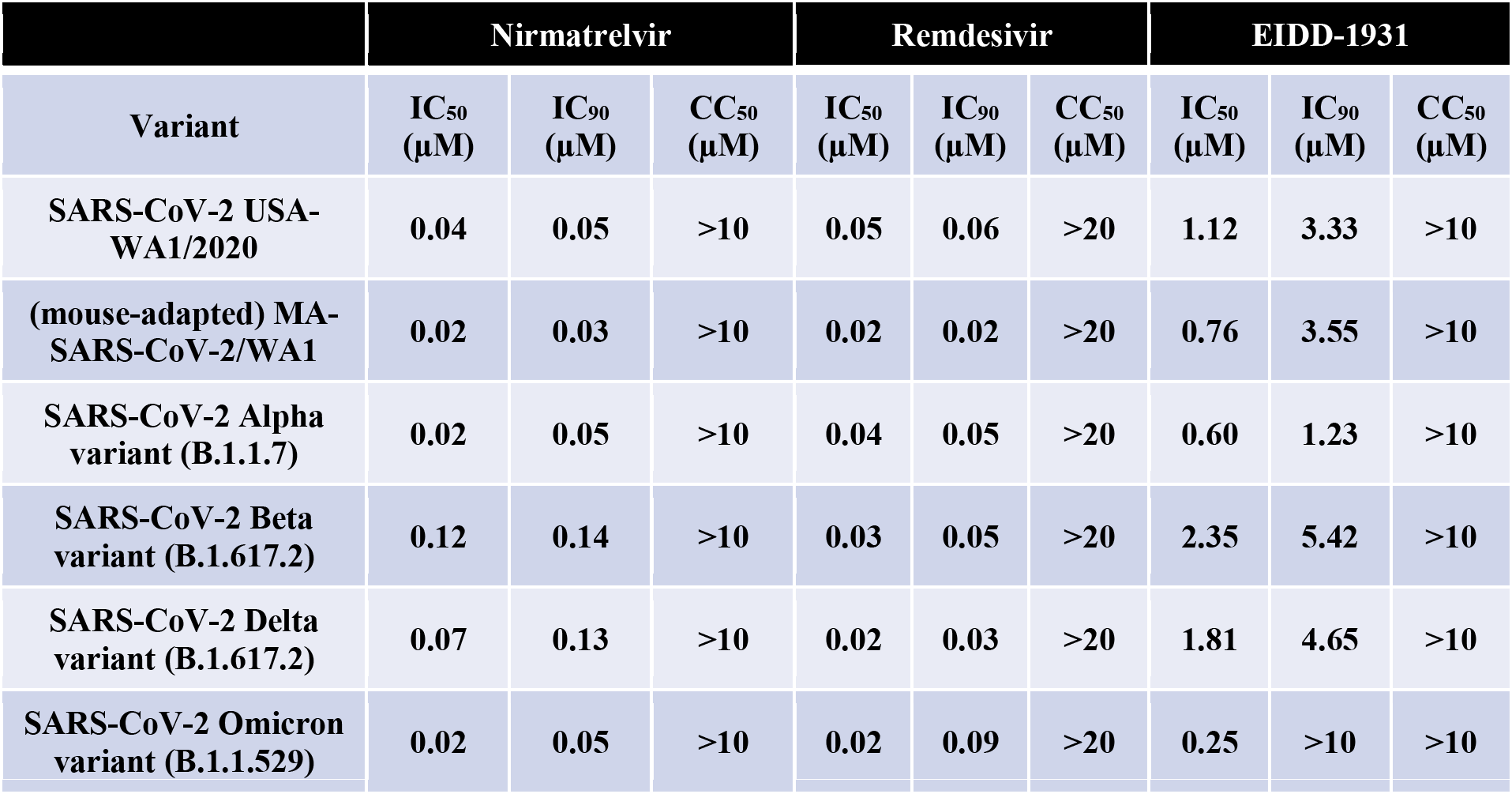
Nirmatrelvir IC_50_, IC_90_, and CC_50_ against clinically relevant SARS-CoV-2 variants in Vero-TMPRSS2 cells (n=3).

We can conclude that nirmatrelvir, molnupiravir, and remdesivir activity against the Delta and Omicron variants, which make up the majority of new clinical infections, is preserved *in vitro*. This aligns with recent observations reported by Vangeel et al.^15^ and Rai et al. (MS ID#: BIORXIV/2022/476644). Therefore, we believe that there is a high probability of maintaining the previously observed clinical activity for nirmatrelvir, molnupiravir, and remdesivir.

These results confirm that therapeutic strategies targeting viral proteins other than the spike protein are robust against many clinically relevant SARS-CoV-2 variants. The SARS-CoV-2 spike protein is under heavy selective pressure, due to being exposed on the surface of the virus and therefore subject to immune pressure. This selective pressure has led to the majority of variant mutations occurring within the spike protein and a subsequent reduction of efficacy for vaccines as antibody titers wane^7,16^, and for most monoclonal antibodies^17,18^. As therapies targeting the SARS-CoV-2 Mpro and RdRp proteins become more widely used, the scientific community will need to closely observe mutations rates within these two viral proteins and monitor whether the deployment of these viral countermeasures impacts the genetic drift towards resistance against nirmatrelvir, remdesivir, and molnupiravir.

## Funding

This study was sponsored by Pfizer Inc. The Mount Sinai Pathogen Surveillance Program is supported, in part, by institutional funds and the SARS-CoV-2 Assessment of Viral Evolution (SAVE) initiative funded through the NIAID Centers of Excellence for Influenza Research and Response (CEIRR) contract 75N93021C00014. Work at the K.M.W. and A.G-S lab on this study was funded by Pfizer; experimental models needed for these studies in the A.G-S lab were possible thanks to the support from CRIPT (Center for Research on Influenza Pathogenesis and Transmission), a NIAID funded Center of Excellence for influenza Research and Response (CEIRR, contract #75N93021C00014), and from NIAID grant U19AI135972, DoD grant W81XWH-20-1-0270 and DARPA grant HR0011-19-2-319 0020.

## Author contributions

R.R., B.L.M., and M.L.R. conducted experiments. R.R., B.L.M., M.L.R, and K.M.W. designed and interpreted data results. R.R., R.D.C., A.S.A, V.S., H.vB., and K.M.W. contributed to writing of manuscript. D.K.R., R.D.C., A.S.A., A.G-S., K.M.W., E.M.S., H.vB., V. S., and the PSP group identified the viral variants of concern isolates and contributed to the methods.

## Acknowledgments

We thank Dr. Randy Albrecht for support with the BSL3 facility and procedures at the ISMMS. The Mount Sinai Pathogen Surveillance Program is grateful to the Molecular Microbiology laboratory team for support and help in conducting timely viral surveillance.

## Competing interests

D.K.R., R.D.C., A.S.A., are employees of Pfizer, were involved with the development of nirmatrelvir, and may own/hold stock options. The K.M.W. and A.G-S. laboratories has received research support from Pfizer, Senhwa Biosciences, Kenall Manufacturing, Avimex, Johnson & Johnson, Dynavax, 7Hills Pharma, Pharmamar, ImmunityBio, Accurius, Nanocomposix, Hexamer, N-fold LLC, Model Medicines, Atea Pharma and Merck. A.G.-S. has consulting agreements for the following companies involving cash and/or stock: Vivaldi Biosciences, Contrafect, 7Hills Pharma, Avimex, Vaxalto, Pagoda, Accurius, Esperovax, Farmak, Applied Biological Laboratories, Pharmamar, Paratus, CureLab Oncology, CureLab Veterinary and Pfizer. A.G.-S. is inventor on patents and patent applications on the use of antivirals and vaccines for the treatment and prevention of virus infections and cancer, owned by the Icahn School of Medicine at Mount Sinai, New York, outside of the reported work.

